# A generic white pupae sex selection phenotype for insect pest control

**DOI:** 10.1101/2020.05.08.076158

**Authors:** CM Ward, RA Aumann, MA Whitehead, K Nikolouli, G Leveque, G Gouvi, E Fung, SJ Reiling, H Djambazian, MA Hughes, S Whiteford, C Caceres-Barrios, TNM Nguyen, A Choo, P Crisp, S Sim, S Geib, F Marec, I Häcker, J Ragoussis, AC Darby, K Bourtzis, SW Baxter, MF Schetelig

## Abstract

Mass releases of sterilized male insects, in the frame of sterile insect technique programs, have helped suppress insect pest populations since the 1950s. In the major horticultural pests *Bactrocera dorsalis, Ceratitis capitata*, and *Zeugodacus cucurbitae*, a key phenotype white pupae (wp) has been used for decades to selectively remove females before releases, yet the gene responsible remained unknown. Here we use classical and modern genetic approaches to identify and functionally characterize causal *wp*^−^ mutations in these distantly related fruit fly species. We find that the wp phenotype is produced by parallel mutations in a single, conserved gene. CRISPR/Cas9-mediated knockout of the *wp* gene leads to the rapid generation of novel white pupae strains in *C. capitata* and *B. tryoni*. The conserved phenotype and independent nature of the *wp*^−^ mutations suggest that this technique can provide a generic approach to produce sexing strains in other major medical and agricultural insect pests.

---

Tephritid species, including the Mediterranean fruit fly (medfly) *Ceratitis capitata*, the oriental fruit fly *Bactrocera dorsalis*, the melon fly *Zeugodacus cucurbitae* and the Queensland fruit fly *Bactrocera tryoni*, are major agricultural pests worldwide^1^. The sterile insect technique (SIT) is a species-specific and environment-friendly approach to control their populations, which has been successfully applied as a component of area-wide integrated pest management programs^2-4^. The efficacy and cost-effectiveness of these large-scale operational SIT applications has been significantly enhanced by the development and use of genetic sexing strains (GSS) for medfly, *B. dorsalis* and *Z. cucurbitae*^5, 6^.

A GSS requires two principal components: a selectable marker, which could be phenotypic or conditionally lethal, and the linkage of the wild type allele of this marker to the male sex, ideally as close as possible to the male determining region. In a GSS, males are heterozygous and phenotypically wild type, whilst females are homozygous for the mutant allele thus facilitating sex separation^6-8^. Pupal color was one of the first phenotypic traits exploited as a selectable marker for the construction of GSS. In all three species, brown is the typical pupae color. However, naturally occurring color mutants such as white pupae (*wp*)^9^ and dark pupae (*dp*)^10^ have occurred in the field or laboratory stocks. The *wp* locus was successfully used as a selectable marker to develop GSS for *C. capitata, B. dorsalis* and *Z. cucurbitae*^6, 11, 12^, however, its genetic basis has never been resolved.

Biochemical studies provided evidence that the white pupae phenotype in medfly is due to a defect in the mechanism responsible for the transfer of catecholamines from the hemolymph to the puparial cuticle^13^. In addition, classical genetic studies showed that the wp phenotype is due to a recessive mutation in an autosomal gene located on chromosome 5 of the medfly genome^9, 14^. The development of translocation lines combined with deletion and transposition mapping and advanced cytogenetic studies allowed the localization of the gene responsible for the wp phenotype on the right arm of chromosome 5, at position 59B of the trichogen polytene chromosome map^15^. In the same series of experiments, the *wp* locus was shown to be tightly linked to a *temperature-sensitive lethal* (*tsl*) gene (position 59B-61C), which is the second selectable marker of the VIENNA 7 and VIENNA 8 GSS currently used in all medfly SIT operational programs worldwide^7, 15^.

The genetic stability of a GSS is a major challenge, mainly due to recombination phenomena taking place between the selectable marker and the translocation breakpoint. To address this risk, a chromosomal inversion called D53 was induced and integrated into the medfly VIENNA 8 GSS (VIENNA 8^D53+^)^6, 8^. Cytogenetic analysis indicated that the D53 inversion spans a large region of chromosome 5 (50B-59C on trichogen polytene chromosome map) with the *wp* locus being inside the inversion, close to its right breakpoint^6^.

Extensive genetic and cytogenetic studies facilitated the development of a physical map of the medfly genome^8, 16^. The annotated gene set provided opportunities for the identification of genes or loci-associated mutant phenotypes, such as the *wp* and *tsl*, used for the construction of GSS^16, 17^. Salivary gland polytene chromosome maps developed for *C. capitata, B. dorsalis, Z. cucurbitae*, and *B. tryoni* show that their homologous chromosomes exhibit similar banding patterns. In addition, *in situ* hybridization analysis of several genes confirmed that there is extensive shared synteny, including the right arm of chromosome 5 where the *C. capitata wp* gene is localized^8^. Interestingly, two recent studies identified SNPs associated with the *wp* phenotype in *C. capitata* and *Z. cucurbitae* that were also on chromosome 5^18, 19^.

In the present study, we employed different strategies involving genetics, cytogenetics, genomics, transcriptomics, gene editing and bioinformatics to identify independent natural mutations in a novel gene responsible for pupal coloration in three tephritid species of major agricultural importance, *C. capitata, B. dorsalis*, and *Z. cucurbitae*. We then functionally characterized causal mutations within this gene in *C. capitata* and *B. tryoni* resulting in development of new white pupae strains. Due to its conserved nature^20^ and widespread occurrence in many insect species of agricultural and medical importance, we also discuss the potential use of this gene as a generic selectable marker for the construction of GSS for SIT applications.

## Results

### Resolving the *B. dorsalis wp* locus with interspecific introgression

The *B. dorsalis* white pupae phenotype was introgressed into *B. tryoni* to generate a strain referred to as the *Bactrocera Introgressed Line* (*BIL*, Supplementary Fig. 1). To determine the proportion of *B. dorsalis* genome introgressed into *BIL*, whole genome sequence data from male and female *B. dorsalis, B. tryoni*, and *BIL* individuals were analyzed. Paired end Illumina short read data from single *B. oleae* males (SRR826808) and females (SRR826807) were used as an outgroup. Single copy orthologs across the genome (n = 1,846) were used to reconstruct the species topology revealing species-specific monophyly (Fig. 1A) consistent with previously published phylogenies^21, 22^. Reconstruction also showed monophyly between *B. tryoni* and *BIL* across 99.2% of gene trees suggesting the majority of loci originally introgressed from *B. dorsalis* have been removed during backcrosses.

**Fig. 1.**
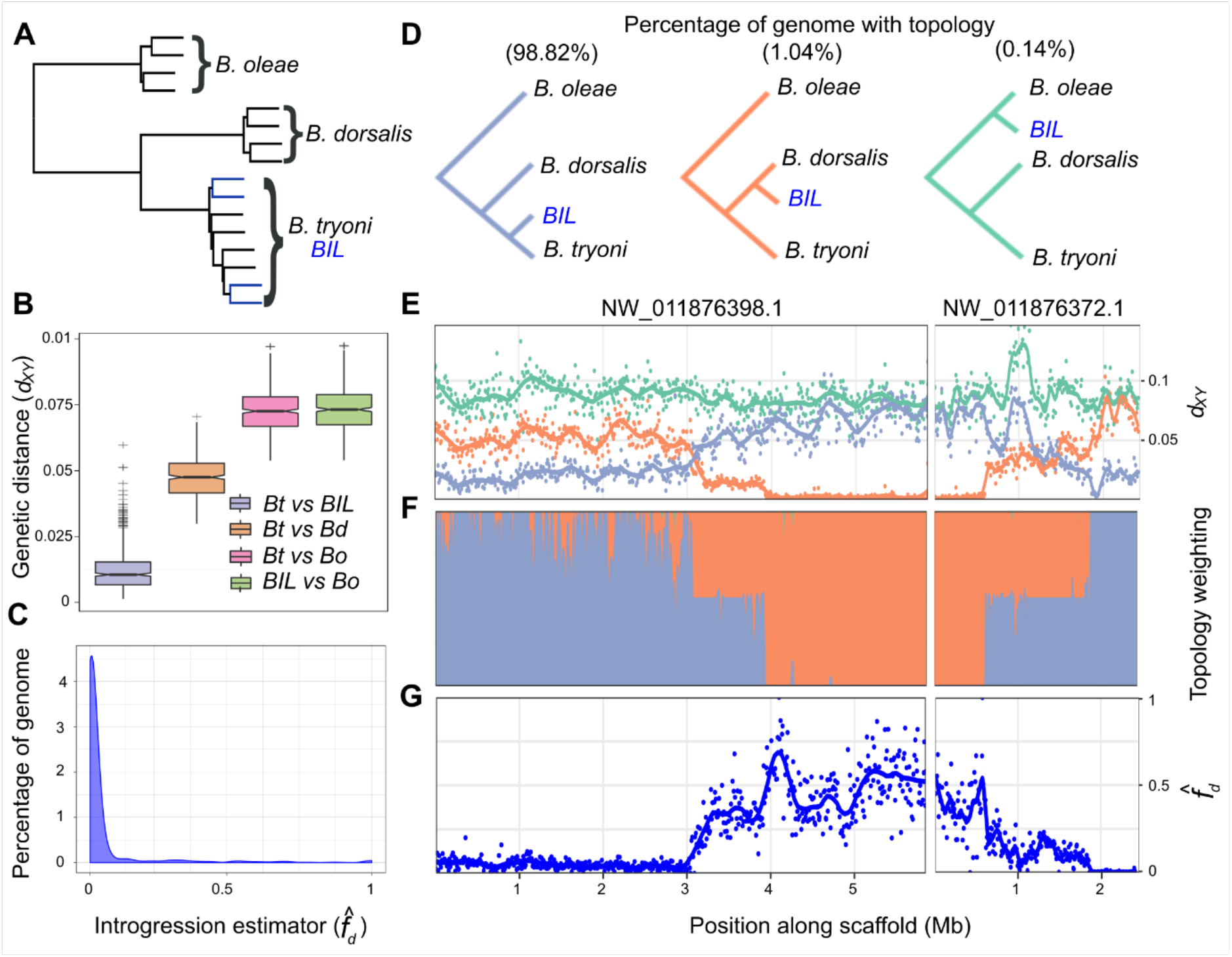
Characterization of total introgression from *B. dorsalis* into the *Bactrocera Introgressed Line* and identification of the *white pupae* locus. (A) Species tree constructed from 1,846 single copy ortholog gene trees for four haplotypes of *B. oleae, B. dorsalis, B. tryoni* and *BIL*. Branches corresponding to *BIL* individuals are shown in blue. All nodes were well supported with posterior probabilities >0.97. (B) Nei’s absolute genetic distance (*d_XY_*) calculated for tiled 100 kb windows across the genome between *B. tryoni* vs *BIL* (*Bt* vs BIL); *B. tryoni* vs *B. dorsalis (Bt* vs *Bd); B. tryoni* vs *B. oleae* (*Bt* vs *Bo*) and *BIL* vs *B. oleae* (*BIL* vs *Bo*). (C) The introgression estimator 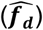 calculated across tiled 100 kb windows to identify regions of disproportionately shared alleles between *BIL* and *B. dorsalis*, 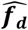 (*Bt, BIL, Bd; Bo*). (D) The three evolutionary hypothesis/topologies of interest to identify introgressed regions and their representation across the genome: species (purple, 98.82%), introgression (orange, 1.04%) and a negative control tree (green, 0.14%). (E) Nei’s absolute genetic distance (*d_XY_*) calculated for tiled 10 kb windows across the candidate *wp* locus colors follow the legend in (D). (F) Topology weighting for each topology shown in (D) calculated for 1 kb tiled local trees across the candidate *wp* locus. (G) The introgression estimator 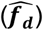 calculated across tiled 10 kb windows, 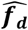 (*Bt, BIL, Bd; Bo*).

Genomes were partitioned into 100 kb windows and pairwise absolute genetic distance (*d_XY_*) calculated between each species and *BIL* to estimate admixture. *B. dorsalis* was found to be highly similar to a small proportion of the *BIL* genome (Fig. 1B; purple), as indicated by *d_XY_* values approaching the median value of *B. dorsalis* vs *B. tryoni* (Fig. 1B; orange).

Two formal tests for introgression were also carried out, the *f* estimator 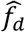 (Fig. 1C) and topology weighting (Fig. 1D). Three distinct local evolutionary histories (Fig. 1D) were tested using *d_XY_* and topology weighting across the *B. dorsalis wp* Quantitative Trait Locus (QTL) i) *BIL* is closest to *B. tryoni* (Fig. 1D; purple, expected across most of the genome), ii) *BIL* is closest to *B. dorsalis* (Fig. 1D; orange, expected at the *wp*^−^ locus), and iii) *BIL* is closest to *B. oleae* (Fig. 1D; green, a negative control). Across the nuclear genome the species topology was supported in 98.82% of windows. Both 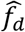 and topology weighting confirmed a lack of widespread introgression from *B. dorsalis* into *BIL* with few (n = 42) discordant outlier windows. Genomic windows discordant across all three tests were considered candidate regions for the *wp* mutation. Four scaffolds accounting for 1.18% of the *B. dorsalis* genome met these criteria and only two, NW_011876372.1 and NW_011876398.1, showed homozygous introgression consistent with a recessive white pupae phenotype (Supplementary Fig. 2).

To resolve breakpoints within the *B. dorsalis wp* QTL, a 10 kb windowed analysis across NW_011876398.1 and NW_011876372.1 was performed using *d_XY_* (Fig. 1E), topology weighting (Fig. 1F) and 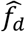 (Fig. 1G). The maximum range of the introgressed locus was 4.49 Mb (NW_011876398.1 was 2.9-5.94 Mb and NW_011876372.1 was 0-1.55 Mb) (Fig. 1E-G). The *wp*^−^ QTL was further reduced to a 2.71 Mb region containing 113 annotated protein coding genes through analyzing nucleotide diversity (*π*) among eight pooled *BIL* genomes (3.8 Mb on NW_011876398.1 to 0.73 Mb on scaffold NW_011876372.1, Supplementary Fig. 2).

### Resolving the *C. capitata* D53 inversion breakpoints and *wp* locus with genome sequencing and *in situ* hybridization

Previous cytogenetic studies determined the localization of the gene responsible for the white pupae phenotype on the right arm of chromosome 5, at position 59B of the trichogen polytene chromosome map^15^. The equivalent of position 59B is position 76B of the salivary gland polytene chromosome map, inside but close to the right breakpoint of the D53 inversion (69C-76B on the salivary gland polytene chromosome map). Long read sequencing data were generated of the wild type strain Egypt II (EgII), the inversion line D53 and the genetic sexing strain VIENNA 8 (without the inversion; VIENNA 8^D53-|-^) (Supplementary Table 1) to enable a comparison of the genomes and locate the breakpoints of the D53 inversion, to subsequently narrow down the target region, and to identify *wp* candidate genes.

Chromosome 5-specific markers^16^ were used to identify the EgII_Ccap3.2.1 scaffold_5 as complete chromosome 5. Candidate D53 breakpoints in EgII scaffold_5 were identified using the alignment of three genome datasets EgII, VIENNA 8^D53-|-^, and D53 (see material and methods). The position of the D53 inversion breakpoints was located between 25,455,334 – 25,455,433 within a scaffold gap (left breakpoint), and at 61,880,224 bp in a scaffolded contig (right breakpoint) on EgII chromosome 5 (Ccap3.2.1; accession GCA_902851445). The region containing the causal *wp* gene was known to be just next to the right breakpoint. Cytogenetic analysis and *in situ* hybridization using the wild type EgII strain and the D53 inversion line confirmed the overall structure of the inversion, covering the area of 69C-76B on the salivary gland polytene chromosomes (Fig. 2), as well as the relative position of markers residing inside and outside the breakpoints (Fig. 2 and Supplementary Fig. 3). PCRs using two primer pairs flanking the predicted breakpoints (Supplementary Fig. 4) and subsequent sequencing confirmed the exact sequence of the breakpoints. Using a primer combination specific for the chromosome 5 wild type status confirmed in the WT in EgII flies and VIENNA 7^D53+|−^ GSS males, which are heterozygous for the inversion. Correspondingly, these amplicons were not present in D53 males and females or in VIENNA 7^D53+|+^ GSS females (all homozygous for the inversion) (Supplementary Fig. 4). Positive signals for the inversion were detected in D53 and VIENNA 7^D53+^ GSS males and females, but not in WT flies using an inversion-specific primer pair (Supplementary Fig. 4).

**Fig. 2.**
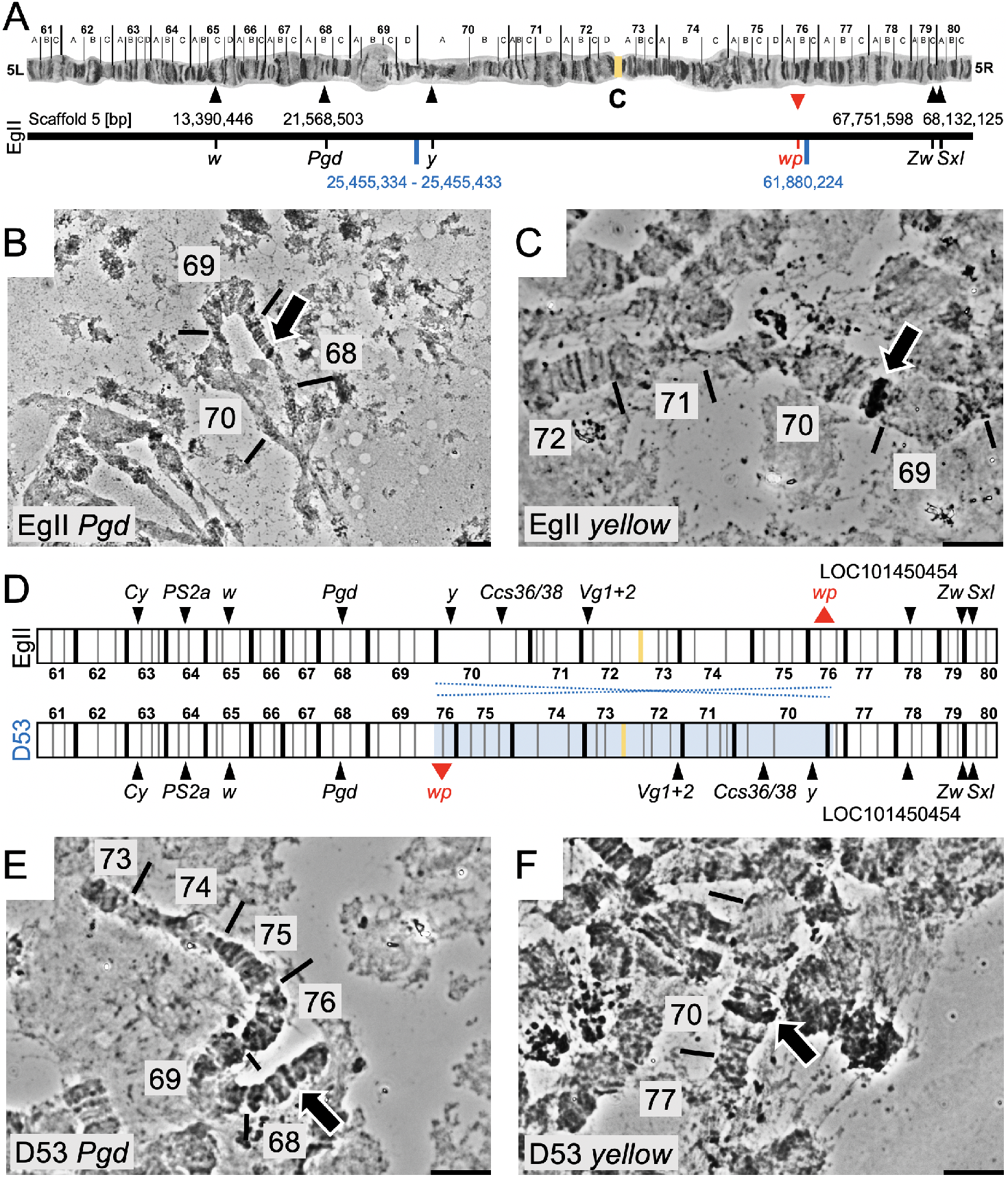
Genomic positioning of the D53 inversion on chromosome 5 of *C. capitata*. (A) Chromosome scale assembly of *C. capitata* EgII chromosome 5. Shown are the positions of *in situ* mapped genes *white* (*w*), *6-phosphogluconate dehydrogenase (Pgd), glucose-6-phosphate 1-dehydrogenase* (*Zw*) and *sex lethal* (*Sxl*), the position of the D53 inversion breakpoints (blue), and the relative position of *white pupae* (*wp*) on the polytene chromosome map of chromosome 5 ^23^ and the PacBio-Hi-C EgII scaffold_5, representing the complete chromosome 5 (genome Ccap3.2.1, accession GCA_902851445). The position of the *yellow* gene (*y*, LOC101455502) was confirmed on chromosome 5 70A by *in situ* hybridization, despite its sequence not been included in the scaffold assembly. (D) Schematic illustration of chromosome 5 without (EgII, WT) and with (D53) D53 inversion. The inverted part of chromosome 5 is shown in light blue. Two probes, one inside (*y*, 70A) and one outside (*Pgd*, 68B) of the left inversion breakpoint were used to verify the D53 inversion breakpoints by *in situ* hybridization. WT EgII is shown in (B) and (C), D53 in (E) and (F).

### Genome and transcriptome sequencing lead to a single candidate *wp* gene in *B. dorsalis, C. capitata, and Z. cucurbitae*

Orthologs within the QTL of *B. dorsalis, C. capitata* and scaffolds previously identified to segregate with the *wp*^−^ phenotype in *Z. cucurbitae* (NW_011863770.1 and NW_011863674.1) ^18^ were investigated for high effect mutations under the assumption that a null mutation in a conserved gene results in the *wp*^−^ phenotype. A single ortholog containing fixed indels absent from wild type strains was identified in each species. White pupae *B. dorsalis* and *BIL* strains showed a 37 bp frame-shift deletion in the first coding exon of LOC105232189 introducing a premature stop codon 210 bp from the transcription start site (Fig. 3A). Presence of the deletion was confirmed *in silico* using whole genome resequencing from the *wp* and wildtype mapped to the reference, and by *de novo* assembly of Illumina RNAseq data transcripts (Fig. 3A).

**Fig. 3.**
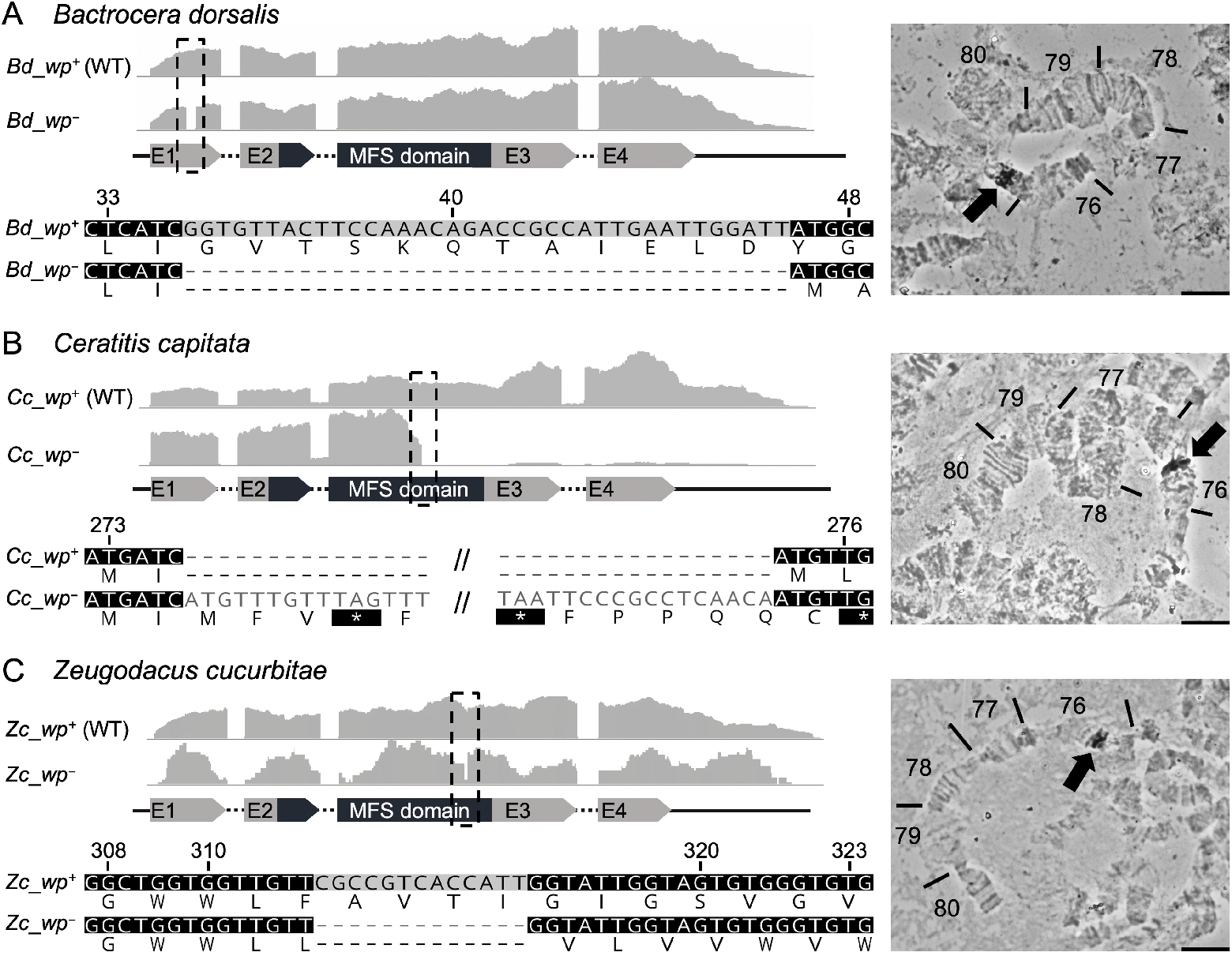
Identification of the *wp* mutation in the transcriptomes of *B. dorsalis, C. capitata* and *Z. cucurbitae*. The grey graphs show expression profiles from the candidate *wp* loci in WT (*wp*^+^) and mutant (*wp*^−^) flies at the immobile pupae stages of (A) *B. dorsalis*, (B) *C. capitata* and (C) *Z. cucurbitae*. The gene structure (not drawn to scale) is indicated below as exons (arrows labelled E1-E4) and introns (dashed lines). The positions of independent *wp* mutations (*Bd*: 37 bp deletion, *Cc*: approximate 8,150 bp insertion, *Zc*: 13 bp deletion) are marked with black boxes in the expression profiles and are shown in detail below the gene models based on *de novo* assembly of RNAseq data from WT and white pupae phenotype individuals. *In situ* hybridization on polytene chromosomes (right) confirmed the presence of the *wp* locus on the right arm of chromosome 5 in all three species (arrows in micrographs).

In *C. capitata, wp* individuals D53 Nanopore read alignment on EgII showed an independent approximate 8,150 bp insertion into the third exon of LOC101451947 disrupting proper gene transcription 822 bp from the transcription start site (Fig. 3B). The insertion sequence is flanked by identical repeats, suggesting that it may originate from a transposable element insertion. The *C. capitata* mutation was confirmed *in silico*, as in *B. dorsalis*, using whole genome sequencing and RNAseq data (Fig. 3B).

Transcriptome data from the white pupae-based genetic sexing strain of *Z. cucurbitae* revealed a 13 bp deletion in the third exon of LOC105216239 on scaffold NW_011863770.1 introducing a premature stop codon (Fig. 3C).

The candidate *white pupae* gene in all three species had a reciprocal best BLAST hit to the putative metabolite transport protein CG14439 in *Drosophila melanogaster* and contains a Major Facilitator-like superfamily domain (MFS_1, pfam07690), suggesting a general function as a metabolite transport protein. *In situ* hybridization on polytene chromosomes of *B. dorsalis, C. capitata* and *Z. cucurbitae* was used to confirm the presence of the *wp* locus in the same syntenic position on the right arm of chromosome 5 (Fig. 3). Therefore, all three species show a mutation in the same positional orthologous gene likely to be responsible for the phenotype in all three genera.

### CRISPR/Cas9 knockout of a Major Facilitator Superfamily gene in *B. tryoni* and *C. capitata* causes the white pupae phenotype

An analogous *B. dorsalis wp*^−^ mutation was developed in *B. tryoni* by functional knockouts of the putative *Bt_wp* using the CRISPR/Cas9 system. A total of 591 embryos from the Ourimbah laboratory strain were injected using two guides with recognition sites in the first coding exon of this gene (Fig. 4A). Injected embryos surviving to adulthood (n = 19, 3.2%) developed with either wild type brown (n = 12) or somatically mosaic white-brown puparia (n = 7, Supplementary Fig. 5). Surviving G_0_ adults were individually backcrossed into the Ourimbah strain, resulting in potentially *wp*^+|−(CRISPR)^ heterozygous brown pupae (Fig. 4C). Five independent G_0_ crosses were fertile (three mosaic white-brown and two brown pupae phenotypes), G_1_ offspring were sibling mated and visual inspection of G_2_ progeny revealed that three families contained white pupae individuals. Four distinct frameshift mutations were observed in screened G_2_ progeny (Fig. 4A) suggesting functional KO of putative *Bt_wp* is sufficient to produce the white pupae phenotype in *B. tryoni*. Capillary sequencing of cloned *Bt_MFS* amplicons revealed deletions ranging from a total of 4 bp to 155 bp, summed across the two guide recognition sites, introducing premature stop codons.

**Fig. 4.**
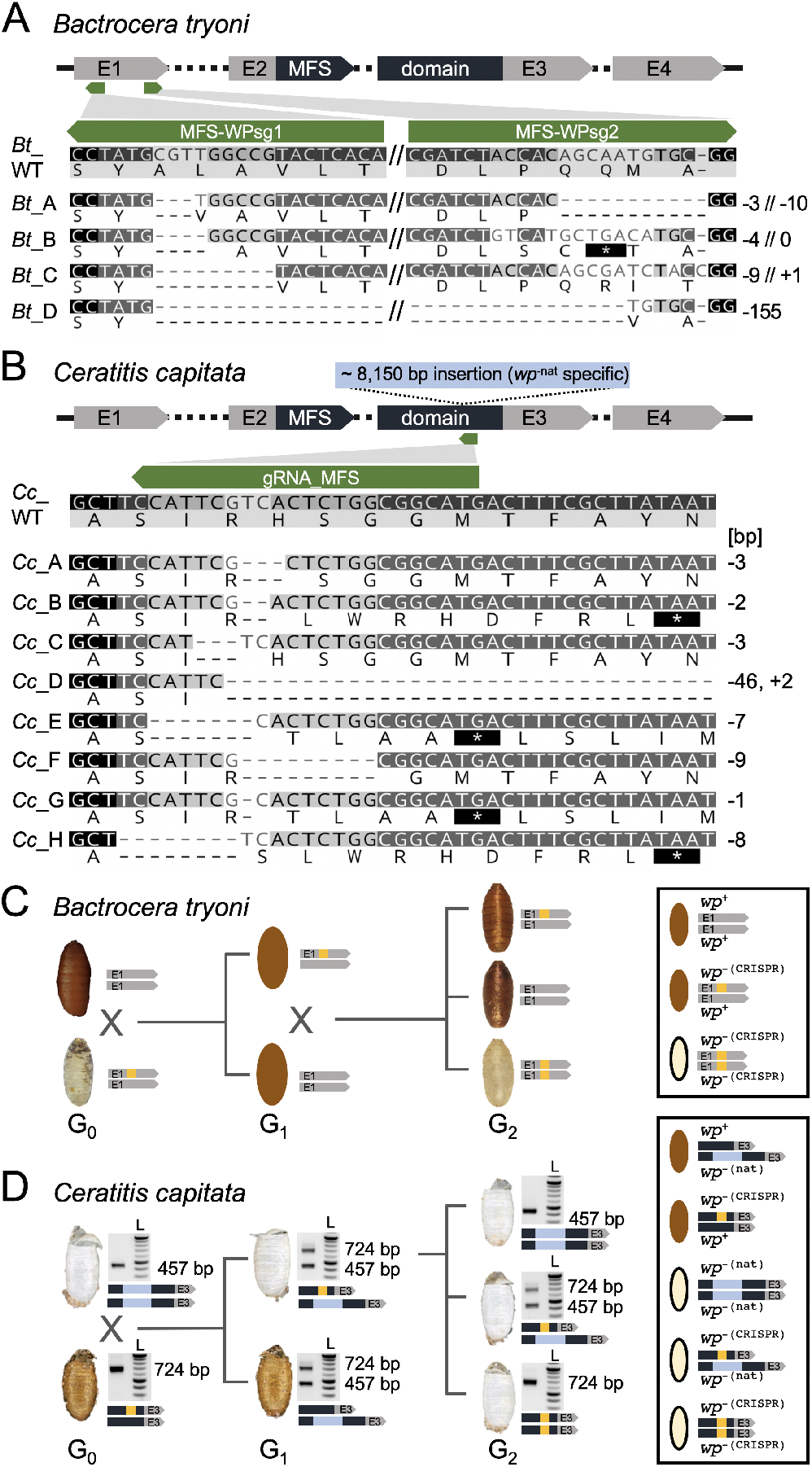
CRISPR/Cas9-based generation of homozygous *wp*^−(CRISPR)^ lines in *B. tryoni* and *C. capitata*. A schematic structure of the *wp* CDS exons (E1, E2, E3, E4) including the MFS domain in *B. tryoni* (A) and *C. capitata* (B) are shown. Positions of gRNAs targeting the first and third exon in *B. tryoni* and *C. capitata*, respectively, are indicated by green arrows. Nucleotide and amino acid sequences of mutant *wp* alleles identified in G_1_ individuals are compared to the WT reference sequence in *B. tryoni* (A) and *C. capitata* (B). Deletions are shown as dashes, alterations on protein level leading to premature stop codons are depicted as asterisks highlighted in black. Numbers on the right side represent InDel sizes (bp = base pairs). Crossing schemes to generate homozygous *wp*^−(CRISPR)^ lines in *B. tryoni* (C) and *C. capitata* (D) show different strategies to generate *wp* strains. Bright field images of empty puparia are depicted for both species. Genotype schematics and corresponding PCR analysis (for *C. capitata*) validating the presence CRISPR-induced (orange) and natural (blue, for *C. capitata) wp* mutations are shown next to the images of the puparia. (C) Injected G_0_ *B. tryoni* were backcrossed to the Ourimbah laboratory strain resulting in uniformly brown G_1_ offspring (depicted as illustration because no images were acquired during G_1_). G_1_ inbreeding led to G_2_ individuals homozygous for the white pupae phenotype. (D) Injected WT G_0_ flies were crossed to flies homozygous for the naturally occurring *wp*^−^ allele (*wp*^−(nat)^). *wp*^−(nat)^ (457 bp amplicon) and *wp*^−(CRISPR)^ or WT (724 bp amplicon) alleles were identified by multiplex PCR (left lane; L = NEB 2log ladder). White pupae phenotypes in G_1_ indicated positive CRISPR events. G_2_ flies with a white pupae phenotype that were homozygous for *wp*-^(CRISPR)^ allele were used to establish lines.

In *C. capitata*, CRISPR/Cas9 gene editing was used to knockout the orthologous gene and putative *Cc_wp*, LOC101451947, to confirm that it causes a *wp*^−^ phenotype. A mix of recombinant Cas9 protein and the gRNA_MFS, targeting the third exon and thereby the MFS domain of the presumed *Cc_wp* CDS (Fig. 4B), was injected into 588 EgII WT embryos of which 96 developed to larvae and 67 pupated. All injected G_0_ pupae showed brown pupal color. In total, 29 G_0_ males and 34 females survived to adulthood (9.3%) and were backcrossed individually or in groups (see material and methods) to a strain carrying the naturally occurring white pupae mutation (*wp*^−(nat)^; strain #1402_22m1B)^24^ (Fig. 4D). As *white pupae* is known to be monogenic and recessive in *C. capitata*, this complementation assay was used to reveal whether the targeted gene is responsible for the naturally occurring white pupae phenotype or if the mutation is located in a different gene. G_1_ offspring would only show white pupae phenotypes if *Cc_wp* was indeed the *white pupae* gene, knocked-out by the CRISPR approach, and complemented by the natural mutation through the backcross (*wp*^−(nat)|-(CRISPR)^). In the case that the *Cc_wp* is not the gene carrying the natural *wp*^−^ mutation, a brown phenotype would be observed for all offspring. Here, five out of 13 crosses, namely M1, M3, F2, F3, and F4, produced white pupae phenotype offspring. The crosses generated 221, 159, 70, 40, and 52 G_1_ pupae, of which 10, 30, 16, 1, and 1 pupa respectively, were white. Fifty-seven flies emerged from white pupae were analyzed via non-lethal genotyping, and all of them showed mutation events within the target region. Overall, eight different mutation events were seen, including deletions ranging from 1-9 bp and a 46 bp deletion combined with a 2 bp insertion (Fig. 4B). Five mutation events (B, D, E, G, H) caused frameshifts and premature stop codons. The remaining three (A, C, F), however, produced deletions of only one to three amino acids. To make *wp*^−(CRISPR)^ mutations homozygous, mutants were either inbred (mutation C) (Fig. 4D) or outcrossed to WT EgII (mutation A-H), both in groups according to their genotype. This demonstrated that *Cc_wp* is the gene carrying the *wp*^−(nat)^, and that even the loss of a single amino acid without a frameshift at this position can cause the white pupae phenotype. Offspring from outcrosses of mutation A, D, and H, as well as offspring of the inbreeding (mutation C), were genotyped via PCR, and *wp*^+|−(CRISPR)^ and *wp*^−(CRISPR)|-(CRISPR)^ positive flies were inbred to establish pure new *white pupae* lines.

## Discussion

*White pupae* (*wp*) was first identified in *C. capitata* as a spontaneous mutation and was subsequently adopted as a phenotypic marker of fundamental importance for the construction of GSS for SIT^6, 9^. Full penetrance expressivity and recessive inheritance rendered *wp* the marker of choice for GSS construction in two additional tephritid species, *B. dorsalis* and *Z. cucurbitae*^11, 12^, allowing automated sex sorting based on pupal color. This was only possible because spontaneous *wp* mutations occur at relatively high rates either in the field or in mass rearing facilities and can easily be detected^6, 9^. Despite the easy detection and establishment of *wp* mutants in these three species, similar mutations have not been detected in other closely or distantly related species such as *B. tryoni, B. oleae*, or *Anastrepha ludens*, despite large screens being conducted. In addition to being a visible GSS marker used to separate males and females, the wp phenotype is also important for detecting and removing recombinants in cases where sex separation is based on a conditional lethal gene such as the *tsl* gene in the medfly VIENNA 7 or VIENNA 8 GSS^6, 7^. However, it took more than 20 years from the discovery and establishment of the *wp* mutants to the large-scale operational use of the medfly VIENNA 8 GSS for SIT applications^6, 9^ and the genetic nature of the *wp* mutation remained unknown. The discovery of the underlying *wp* mutations and the availability of CRISPR/Cas genome editing would allow the fast recreation of such phenotypes and sexing strains in other insect pests. Isolation of the *wp* gene would also facilitate future efforts towards the identification of the closely linked *tsl* gene.

Using an integrated approach consisting of genetics, cytogenetics, genomics, transcriptomics and bioinformatics, we identified the white pupae (*wp*) genetic locus in three major tephritid agricultural pest species, *B. dorsalis, C. capitata*, and *Z. cucurbitae*.

Our study clearly shows the power of employing different strategies for gene discovery, one of which was species hybridization. In *Drosophila*, hybridization of different species has played a catalytic role in the deep understanding of species boundaries and the speciation processes, including the evolution of mating behavior and gene regulation^25-29^. In our study, we took advantage of two closely related species, *B. dorsalis* and *B. tryoni*, which can produce fertile hybrids and be backcrossed for consecutive generations. This allowed the introgression of the *wp* mutant locus of *B. dorsalis* into *B. tryoni*, resulting in the identification of the introgressed region, including the causal *wp* mutation via whole-genome resequencing and advanced bioinformatic analysis.

In *C. capitata*, we exploited two essential pieces of evidence originating from previous genetic and cytogenetic studies: the localization of *wp* to region 59B and 76B on chromosome 5 in the trichogen cells and salivary gland polytene chromosome map, respectively^15, 30^, and its position close to the right breakpoint of the large inversion D53^6^. This data prompted us to undertake a comparative genomic approach to identify the exact position of the right breakpoint of the D53 inversion, which would bring us in the vicinity of the *wp* gene. Coupled with comparative transcriptomic analysis, this strategy ensured that the analysis indeed tracked the specific *wp* locus on the right arm of chromosome 5, instead of any mutation in another, random locus which may participate in the pigmentation pathway and therefore result in the same phenotype. Functional characterization via CRISPR/Cas9-mediated knockout resulted in the establishment of new white pupae strains in *C. capitata* and *B. tryoni* and confirmed that this gene is responsible for the pupal coloration in these tephritid species. Interestingly, the wp phenotype is based on three independent and very different natural mutations of this gene, a rather large and transposon-like insertion in *C. capitata*, but only small deletions in the two other tephritids, *B. dorsalis* and *Z. cucurbitae*. In medfly, however, CRISPR-induced in-frame deletions of one or three amino acids in the MFS domain were sufficient to induce the wp phenotype, underlining the importance of this domain for correct pupal coloration.

It is worth noting, that in the first stages of this study, we employed two additional approaches, which did not allow us to successfully narrow down the *wp* genomic region to the desired level. The first was based on Illumina sequencing of libraries produced from laser micro-dissected (Y;5) mitotic chromosomes that carry the wild type allele of the *wp* gene through a translocation from the fifth chromosome to the Y. This dataset from the medfly VIENNA 7 GSS was comparatively analyzed to wild type (Egypt II) Y and X chromosomes, and the complete genomes of Egypt II, VIENNA 7 ^D53-^ GSS, and a D53 inversion line in an attempt to identify the chromosomal breakpoints of the translocation and/or inversion, which are close to the *wp* locus (Supplementary Table 2). However, this effort was not successful due to the short Illumina reads and the lack of a high-quality reference genome. The second approach was based on individual scale whole-genome resequencing/genotyping, and identifying fixed loci associated with pupal color phenotypes, which complemented the QTL analysis^19^. Seven loci associated with SNPs and larger deletions linked to the white pupae phenotype were analyzed based on their respective mutations and literature searches for their potential involvement in pigmentation pathways. However, we could not identify a clear link to the pupal coloration as shown by *in silico*, molecular, and *in situ* hybridization analysis (Supplementary Fig. 6 and 7, Supplementary Table 3).

The *wp* gene is a member of a major facilitator superfamily (MFS). Orthologs of *white pupae* are present in 146 of 148 insect species aggregated in OrthoDB v9^20^ present in all orders and single copy in 133 species. Furthermore, *wp* is included in the benchmarking universal single copy ortholog (BUSCO) gene set for insecta and according to OrthoDB v10^31^ has a below average evolutionary rate (0.87, OrthoDB group 42284at50557) suggesting an important and evolutionarily conserved function (Supplementary Fig. 8). Its ortholog in *Bombyx mori, mucK*, was shown to participate in the pigmentation at the larval stage^32^ whereas in *D. melanogaster* peak expression is during the prepupal stage after the larva has committed to pupation^33^, which is the stage where pupal cuticle sclerotization and melanization occurs. It is known that the insect cuticle consists of chitin, proteins, lipids and catecholamines, which act as cross-linking agents thus contributing to polymerization and the formation of the integument^34^. Interestingly, the sclerotization and melanization pathways are connected and this explains the different mechanical properties observed in different medfly pupal color strains with the “dark” color cuticles to be harder than the “brown” ones and the latter harder than the “white” color ones^35^. The fact that the white pupae mutants are unable to transfer catecholamines from the haemolymph to the cuticle is perhaps an explanation for the lack of the brown pigmentation^13^.

The discovery of the long-sought *wp* gene in this study and the recent discovery of the *Maleness-on-the-Y (MoY*) gene, which determines the male sex in several tephritids^36^, opens the way for the development of a ‘generic approach’ for the construction of GSS for other species. Using CRISPR/Cas-based genome editing approaches, we can: (a) induce mutations in the *wp* orthologues of SIT target species and establish lines with wp phenotype and (b) link the rescue alleles as closely as possible to the *MoY* region. Given that the *wp* gene is present in diverse insect species including agricultural insect pests and mosquito disease vectors, this approach would allow more rapid development of GSS in SIT target species. In principle, these GSS will have higher fertility compared to the semi-sterile translocation lines^6^. In addition, these new generation GSS will be more stable since the rescue allele will be tightly linked to the male determining region thus eliminating recombination which can jeopardize the genetic integrity of any GSS. The concept of the ‘generic approach’ can also be applied in species which lack a typical Y chromosome such as *Aedes aegypti* and *Aedes albopictus*. In these species, the rescue allele should be transferred close to the male determining gene (*Nix*) and the M locus^37, 38^. It is hence important for this ‘generic approach’ to identify regions close enough to the male determining loci to ensure the genetic stability of the GSS and to allow the proper expression of the rescue alleles. In the present study, we have already shown that CRISPR/Cas9-induced mutations resulting in the white pupae phenotype can be developed in SIT target species and the resulting strains provide already new opportunities for GSS based on visible markers.

## Methods

All sequence libraries prepared during this study are publicly available on NCBI within the ENA BioProject PRJEB36344/ERP119522 (accession numbers ERS4426857 – ERS4426873, ERS4426994 – ERS4427029, ERS4519515, ERS4547590 – ERS4547593; see Supplementary Table 1) and the BioProject PRJNA629430 (SRA accessions SRR11649127 – SRR11649132; see Supplementary Fig. 6).

### Insect rearing

*C. capitata, B. dorsalis* and *Z. cucurbitae* fly strains were maintained at 25±1°C, 48% RH and 14/10 h light/dark cycle. They were fed with a mixture of sugar and yeast extract (3v:1v) and water. Larvae were reared on a gel diet, containing carrot powder (120 g/l), agar (3 g/l), yeast extract (42 g/l), benzoic acid (4 g/l), HCl (25%, 5.75 ml/l) and ethyl-4- hydroxybenzoate (2.86 g/l). Flies were anesthetized with N_2_ or CO_2_ for screening, sexing, and the setup of crosses. To slow down the development during the non-lethal genotyping process (*C. capitata*), adult flies were kept at 19°C, 60% RH, and 24 h light for this period (1-4 d).

*B. tryoni* flies were obtained from New South Wales Department of Primary Industries (NSW DPI), Ourimbah, Australia and reared at 25 ± 2°C, 65 ± 10% RH and 14/10 h light/dark cycle. Flies were fed with sugar, Brewer’s yeast and water and larvae were reared on a gel diet, containing Brewer’s yeast (204 g/l), sugar (121 g/l), methyl p-hydroxy benzoate (2 g/l), citric acid (23 g/l), wheat germ oil (2 g/l), sodium benzoate (2 g/l) and agar (10 g/l).

### Introgression and bioinformatic identification of a natural *wp* mutation from *B. dorsalis* in *B. tryoni*

Interspecific crosses between *B. tryoni* and *white pupae B. dorsalis* were carried out between male *B. tryoni* (*wp*^+|+^) and female *B. dorsalis* (*wp*^−|-^). The G_1_ *wp*^+|−^ hybrids developed with brown puparia and were mass crossed. G_2_ *wp*^−|-^ females were backcrossed into *B. tryoni wp*^11^ males and backcrossing was then repeated for five additional times to produce the *white pupae Bactrocera Introgressed Line (BIL*, Supplementary Fig. 1).

Genome sequencing using Illumina NovaSeq (2 × 150 bp, Deakin University) was performed on a single male and female from the *B. dorsalis wp* strain, *B. tryoni*, and the *BIL* (~ 26X) and two pools of five *BIL* individuals (~32X). Quality control of each sequenced library was carried out using FastQC v0.11.6 (https://www.bioinformatics.babraham.ac.uk/projects/fastqc/) and aggregated using ngsReports^39^ v1.3. Adapter trimming was carried out using Trimmomatic v0.38 (https://academic.oup.com/bioinformatics/article/30/15/2114/2390096) and paired reads were mapped to the *B. dorsalis* reference genome (GCF_000789215.1) using NextGenMap^40^ v0.5.5 under default settings. Mapped data were sorted and indexed using SAMtools, and deduplication was carried out using Picard MarkDuplicates v2.2.4 (https://github.com/broadinstitute/picard). Genotypes were called on single and pooled libraries separately with ploidy set to two and ten respectively using Freebayes^41^ v1.0.2. Each strain was set as a different population in Freebayes. Genotypes with less than five genotype depth were set to missing and sites with greater than 20% missing genotypes or indels filtered out using BCFtools^42^ v1.9. Conversion to the genomic data structure (GDS) format was carried out using SeqArray^43^ v1.26.2 and imported into the R package geaR v0.1 (https://github.com/CMWbio/geaR) for population genetic analysis.

Single copy orthologs were identified in the *B. dorsalis* reference annotated proteins (NCBI *Bactrocera dorsalis* Annotation Release 100) with BUSCO^44, 45^ v3 using the dipteran gene set^44^. Nucleotide alignments of each complete single copy ortholog were extracted from the called genotype set using geaR v0.1 and gene trees built using RAxML^46^ v8.2.10 with a GTR+G model. Gene trees were then imported into Astral III v5.1.1 ^47^ for species tree estimation. Genome scans of absolute genetic divergence (d_XY_), nucleotide diversity (π), and the f estimator 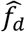 were carried out using geaR v0.1. Two levels of analysis were carried out: i) genome wide scans of non-overlapping 100 kb windows and ii) locus scans of 10 kb tiled windows. Local phylogenies were built for nucleotide alignments of non-overlapping 1 kb windows using RAxML v8.2.10 with a GTR+G model and topology weighting was calculated using TWISST^48^.

Introgressed regions (i.e. candidate *wp* loci) were identified by extracting windows in the genome wide scan with topology weighting and fd greater than 0.75 and visually inspecting the ‘locus scan’ data set for d_XY_, fd and topology weighting patterns indicative of introgression. Nucleotide alignments of all genes within candidate *B. dorsalis* introgressed regions were extracted from the GDS using geaR and visually inspected for fixed mutations in *B. dorsalis wp*, BIL individuals and the two BIL pools. Candidate genes were then searched by tBLASTn against the *D. melanogaster* annotated protein set to identify putative functions and functional domains were annotated using HMMer^49^. Mapped read depth was calculated around candidate regions using SAMtools^50^ depth v1.9 and each sample’s read depth was normalized to the sample maximum to inspect putative deletions. Called genotypes were confirmed by *de novo* genome assembly of the *B. dorsalis wp* genome using MaSuRCA^51^ v3.3 under default settings. The *de novo* scaffold containing LOC105232189 was identified using the BLASTn algorithm. *In silico* exon-intron boundaries were then manually annotated in Geneious^52^ v11.

### Identification, characterization and molecular analysis of the D53 inversion and *wp* locus in *C. capitata*

#### Characteristics of *C. capitata* strains used for this study

Egypt II (EgII) is a wild type laboratory strain. D53 is a homozygous strain with an irradiation-induced inversion covering the area 69C-76B on the salivary gland polytene chromosome map (50B-59C on the trichogen cells polytene chromosome map). VIENNA 7 and VIENNA 8 are two GSS in which two (Y;5) translocations, on the region 58B and 52B of the trichogen cells polytene chromosome map respectively, have resulted to the linkage of the wild type allele of the *wp* and *tsl* genes to the male determining region of the Y chromosome. Thus, VIENNA 7 and VIENNA 8 males are heterozygous in the *wp* and *tsl* loci but phenotypically wild type while VIENNA 7 and VIENNA 8 females are homozygous for the mutant alleles and phenotypically white pupae and they die when exposed at elevated temperatures. The VIENNA 7 and VIENNA 8 GSS can be constructed with and without the D53 inversion (VIENNA 7/8 ^D53+^ or ^D53-^). When the GSS have the inversion, females are homozygous (^D53+|+^) for D53 while males are heterozygous (^D53+|−^)^6, 8, 16^.

#### Whole genome sequencing of *C. capitata* strains

High-molecular-weight (HMW) DNA was extracted from *C. capitata* lines (males and females of the wild type EgII strain, the VIENNA 7 ^D53-|-^ and VIENNA 8 ^D53-|-^ GSS and the inversion line D53) and sequenced. Freshly emerged, virgin and unfed males and females were collected from all strains. For 10X Genomics linked read and Nanopore sequencing, the HMW was prepared as follows: twenty individuals of each sex and strain were pooled, ground in liquid nitrogen, and HMW DNA was extracted using the QIAGEN Genomic tip 100/G kit (Qiagen, Germany). For PacBio Sequel an EgII line was created with single pair crossing and subsequent sibling-mating for six generations. In all generations adult and larval diet contained 100 μg/ml tetracycline. HMW DNA from G6 individuals was prepared as follows: five males from this EgII line were pooled and ground in liquid nitrogen, and HMW DNA was extracted using the phenol/chloroform Phase Lock Gel™ tubes (QuantaBio)^53^. DNA for Illumina applications was extracted from individual flies (Supplementary Table 1).

PacBio *de novo* sequencing: samples were purified with AMPure beads (Beckman Coulter, UK) (0.6 volumes) and QC checked for concentration, size, integrity and purity using Qubit (Qiagen, UK), Fragment Analyser (Agilent Technologies) and Nanodrop (Thermo Fisher) machines. The samples were then processed without shearing using the PacBio Express kit 1 for library construction and an input of 4 μg DNA following the manufacturer’s protocol. The final library was size-selected using the Sage Blue Pippin (Sage Sciences) 0.75% cassette U1 marker in the range of 25-80 kb. The final library size and concentrations were obtained on the Fragment Analyser before being sequenced using the Sequel 1 2.1 chemistry with V4 primers at a loading on plate concentration of 6 pM and 10 h movie times.

For Nanopore sequencing, the ligation sequencing kits SQK-LSK109 or SQK-RAD004 were used as recommended by the manufacturer (Oxford Nanopore Technologies, Oxford, United Kingdom). Starting material for the ligation library preparation were 1 – 1.5 μg HMW gDNA for the ligation libraries and 400 ng for the rapid libraries. The prepared libraries were loaded onto FLO-PRO002 (R9.4) flow cells. Data collection was carried out using a PromethION Beta with live high accuracy base calling for up to 72 h and with mux scan intervals of 1.5 h. Each sample was sequenced at least twice. Data generated were 7.7 Gb for EgII male, 31.09 Gb for D53 male, 26.72 Gb for VIENNA 7^D53-|-^ male, and 24.83 Gb for VIENNA 8 ^D53-|-^ male. Run metrics are shown in Supplementary Table 5.

The PacBio data were assembled using CANU^54^ with two parameter settings: the first to avoid haplotype collapsing (genomeSize=500m corOutCoverage=200 “batOptions=-dg 3 -db 3 -dr 1 -ca 500 -cp 50”) and the second to merge haplotypes together (genomeSize=500m corOutCoverage=200 correctedErrorRate=0.15). The genome completeness was assessed with BUSCO^44, 45^ v3 using the dipteran gene set^44^. The two assemblies were found to be “duplicated” due to alternative haplotypes. To improve the contiguity and reduce duplication haploMerger2 was used^55^. The new assembly was then retested with BUSCO v3 and used scaffolding. Phase Genomics™ Hi-C libraries were made by Phase genomics from males (n=2) of the same family used for PacBio sequencing. Initial scaffolding was completed by Phase Genomics, but edited using the Salsa^56^ and 3D-DNA (https://github.com/theaidenlab/3d-dna) software. The resulting scaffolds were allocated a chromosome number using chromosome specific marker described previously^16^. Specific attention was made to the assembly and scaffolding of chromosome 5. Two contig misassemblies were detected by the Hi-C data and fitted manually. The new assembly (EgII_Ccap3.2.1) was then validated using the Hi-C data. Genes were called using the Funannotate software making use of the Illumina RNAseq data generated by this project; mRNA mapping to the genome is described below.

#### D53 breakpoint analysis in *C. capitata*

To identify possible breakpoint positions, the Nanopore D53 fly assembly contig_531 was mapped onto the EgII_scaffold_5 (from the EgII_CCAP3.2_CANU_Hi-C_scaffolds.fasta assembly) using MashMap v2.0 (https://github.com/marbl/MashMap). This helped to visualize the local alignment boundaries (Supplementary Fig. 9). MashMap parameters were set to kmer size = 16; window size = 100; segment length = 500; alphabet = DNA; percentage identity threshold = 95%; filter mode = one-to-one.

Subsequent to this, and to help confirm the exact location of the identified breakpoints, minimap2 (v2.17, https://github.com/lh3/minimap2) was used to align D53 as well as VIENNA 8^D53-|-^ and VIENNA 7^D53-|-^ Nanopore reads onto the EgII scaffold_5 reference (Supplementary Fig. 9). Minimap2 parameters for Nanopore reads were: minimap2 -x map-ont -A 1 -a --MD – L -t 40. Samtools (v1.9, https://github.com/samtools/samtools) was used to convert the alignment.sam to .bam and prepare the alignment file to be viewed in the Integrative Genomics Viewer (IGV, http://software.broadinstitute.org/software/igv/). The expectation was to see a leftmost breakpoint in D53 read set alignments but not in VIENNA 8^D53-|-^ and VIENNA 7^D53-|-^ when compared to the EgII reference (Supplementary Fig. 9). Due to an assembly gap in the EgII scaffold_5 sequence, the exact location of the leftmost inversion breakpoint was not conclusive using this approach.

A complementary approach was then used to facilitate detection of the leftmost inversion breakpoint in the D53 inversion line. Minimap2 was again used, but here D53 contig_531 was used as reference for the mapping of EgII male PacBio reads as well as VIENNA ^8D53-|-^ male and VIENNA 7 D53-|- male Nanopore reads (Supplementary Fig. 10). Minimap2 parameters for PacBio reads were: minimap2 -x map-pb -A 1 -a --MD -L -t 40. Minimap2 parameters for Nanopore reads were: minimap2 -x map-ont -A 1 -a --MD -L -t 40. Samtools (v1.9, https://github.com/samtools/samtools) was used to convert the alignment .sam to .bam and prepare the alignment file to be viewed in the Integrative Genomics Viewer (IGV, http://software.broadinstitute.org/software/igv/). The expectation was to see a common breakpoint for all three of the above read set alignments when compared to the D53 genome in the area of the inversion. Position ~3,055,294 was identified in the D53 contig_531 as the most likely leftmost breakpoint.

To determine the rightmost breakpoint, D53 male and VIENNA 8 ^D53-|-^ and VIENNA 7^D53-|-^ male nanopore reads were aligned on the EgII_scaffold_5 sequence. The expectation was to see a breakpoint in D53 read set alignments but not in VIENNA 7^D53-|-^ and VIENNA 8 ^D53-|-^. This is the case here, since read alignments coming from both sides of the inversion are truncated at one position (Supplementary Fig. 9). Findings from genome version EgII_Ccap3.2 were extrapolated to the manually revised genome version EgII_Ccap3.2.1.

Predicted D53 inversion breakpoints were verified via PCRs in EgII, D53, and VIENNA 7^D53+^ GSS male and female flies, using PhusionFlash Polymerase in a 10 μl reaction volume [98°C, 10 s; 30 cycles of (98°C, 1 s; 56°C, 5 s; 72°C, 35 s); 72°C, 1 min] (Supplementary Fig. 4). The primer pair for the right breakpoint was designed based on EgII sequence information, primers for the left breakpoint were designed based on D53 sequence information. The wild type status of chromosome 5 (EgII male and female, VIENNA 7 ^D53+|−^ male) was amplified using primer pairs P_1794 and P_1798 (1,950 bp) and P_1795 and P_1777 (690 bp) (Supplementary Table 4). Chromosome 5 with the inversion (D53 male and female, VIENNA 7 ^D53+|−^ male and VIENNA 7 ^D53+|+^ female) was verified using primer pairs P_1777 and P_1798 (1,188 bp) and P_1794 and P_1795 (1,152 bp) and amplicon sequencing (Macrogen Europe, Amsterdam).

#### RNA extraction for transcriptomic analysis of *C. capitata, B. dorsalis* and *Z. cucurbitae*

Samples of *C. capitata, B. dorsalis* and *Z. cucurbitae* species were collected for RNA extraction (Supplementary Table 1) at 3rd instar larval and pre-pupal stages. Total RNA was extracted by homogenizing three larvae of *C. capitata* and *B. dorsalis* and a single larvae of *Z. cucurbitae* in liquid nitrogen, and then using the RNeasy Mini kit (Qiagen). Three replicates per strain and time point were performed. mRNA was isolated using the NEBNext polyA selection and the Ultra II directional RNA library preparation protocols from NEB and sequenced on the Illumina NovaSeq 6000 using duel indexes as 150 bp paired end reads (library insert 500 bp). Individual libraries were sequenced to provide >1 million paired end reads per sample. Each replicate was then assembled separately using Trinity^57^. The assembled transcripts from Trinity were mapped to the Ccap3.2 genome using minimap^58^ (parameters -ax splice:hq -uf). The Illumina reads were mapped with STAR^59^. IGV^60^ v 2.6 was used to view all data at a genomic and gene level. Given that the white pupae GSS^12, 61^ was used to collect samples for RNA extraction from single larvae of *Z. cucurbitae*, larval sex was confirmed by a maleness-specific PCR on the *MoY* gene of *Z. cucurbitae*^36^ using cDNA synthesized with the OneStep RT-PCR Kit (Qiagen) and the primer pair ZcMoY1F and ZcMoY1R (Supplementary Table 4) amplifying a 214 bp fragment. Conditions for a 25 μl PCR reaction using the 1 × *Taq* PCR Master Mix kit (Qiagen) were: [95°C, 5 min; 30 cycles of (95°C, 1 min; 51 °C, 1 min; 72°C, 1 min); 72°C, 10 min]. Presence of a PCR product indicated a male sample. Each, male and female sample was a pool of three individuals. Three replicates per strain and time point were collected.

### Functional and cytogenetic verification of medfly D53 inversion and tephritid *wp* genes

Polytene chromosomes for *in situ* hybridization were prepared from third-instar larvae salivary glands as described previously^62^. In brief, the glands were dissected in 45% acetic acid and placed on a coverslip in a drop of 3:2:1 solution (3 parts glacial acetic acid: 2 parts water: 1- part lactic acid) until been transparent (approximately 5 min). The coverslip was picked up with a clean slide. After squashing, the quality of the preparation was checked by phase contrast microscope. Satisfactory preparations were left to flatten overnight at −20°C and dipped into liquid nitrogen until the babbling stopped. The coverslip was immediately removed with razor blade and the slides were dehydrated in absolute ethanol, air dried and kept at room temperature.

Probes were prepared by PCR. Single adult flies were used to extract DNA with the Extract me kit (Blirt SA), following the manufacturer’s protocol. NanoDrop spectrometer was used to assess the quantity and quality of the extracted DNA which was then stored at −20°C until used. Primers (P1790/1791, P1821/1822, Pgd_probe_F/R, vg1_probe_F/R, Sxl_probe_F/R, y_probe_F/R, zw_probe_F/R, P1633/1634, Zc_F/R, Bd_F/R, P1395/1396, P1415/1416; Supplementary Table 4) were designed for each targeted gene using Primer 3 and/or Geneious Prime programs. PCR was performed in a 25 μl reaction volume using 12.5 μl PCR Master mix 2x Kit (Thermo Fisher Scientific), 60-80 ng DNA, and the following PCR settings [94°C, 5 min; 35 cycles of (94°C, 45 s; 56°C, 30 s; 72°C, 90 s); 72°C, 1 min].

The labelling of the probes was carried out according to the instruction manual of the Dig DNA labelling kit (Roche). *In situ* hybridization was performed as described previously^63^. In brief, before hybridization, stored chromosome preparations were hydrated by placing them for 2 min at each of the following ethanol solutions: 70%, 50% and 30%. Then they were placed in 2× SSC at room temperature for 2 min. The stabilization of the chromosomes was done by placing them in 2× SSC at 65°C for 30 min, denaturing in 0.07 M NaOH 2 min, washing in 2× SSC for 30 sec, dehydrating (2 min in 30%, 50%, 70% and 95% ethanol), and air drying. Hybridization was performed on the same day by adding 15 μl of denatured probe (boiled for 10 min and ice-chilled). Slides were covered with a siliconized coverslip, sealed with rubber cement, and incubated at 45°C overnight in a humid box. At the end of incubation, the coverslip was floated off in 2x SSC and the slide washed in 2x SSC for 3 × 20 min at 53°C.

After 5 min wash in Buffer 1 (100 mM tris-HCl pH 7.5/ 1.5 M NaCl), the preparations were in Blocking solution (Blocking reagent 0.5% in Buffer 1) for 30 min, and then washed for 1 min in Buffer 1. The antibody mix was added to each slide and a coverslip was added. Then the slides were incubated in a humid box for 45 min at room temperature, following 2× 15 min washes in Buffer 1, and a 2 min wash in detection buffer (100 mM Tris-HCl pH 9.5/ 100 mM NaCl). The color was developed with 1 ml of NBT/BCIP solution during a 40 min incubation in the dark at room temperature. The removal of the NBT/BCIP solution was done by rinsing in water twice. Hybridization sites were identified using 40x or 100x oil objectives (phase or bright field) and a Leica DM 2000 LED microscope, with reference to the salivary gland chromosome maps^64^. Well-spread nuclei or isolated chromosomes were photographed using a digital camera (Leica DMC 5400) and the LAS X software. All *in situ* hybridizations were performed at least in duplicates and at least 10 nuclei were analyzed per sample.

### Gene editing and generation of homozygous *wp*^−^ strains in *B. tryoni* and *C. capitata*

For CRISPR/Cas9 gene editing in *B. tryoni*, purified Cas9 protein (Alt-R S.p. Cas9 Nuclease V3, #1081058, 10 μg/μl) and guide RNAs (customized Alt-R^®^ CRISPR/Cas9 crRNA, 2 nmol and Alt-R CRISPR/Cas9 tracrRNA, #1072532, 5nmol) were obtained from Integrated DNA Technologies (IDT) and stored at −20°C. The guide RNAs were individually resuspended to a 100 μM stock solution with nuclease-free duplex buffer and stored at −20°C before use. The two customized 20 bp crRNA sequences (Bt_MFS-1 and Bt_MFS-2) (Supplementary Table 4) were designed using CRISPOR^65^. Injection mixes for microinjection of *B. tryoni* embryos comprise of 300 ng/μl Cas9 protein, 59 ng/μl of each individual crRNA, 222 ng/μl tracrRNA and 1x injection buffer (0.1 mM sodium phosphate buffer pH 6.8, 5 mM KCI) in a final volume of 10 μl. The guide RNA complex containing the two crRNAs and tracrRNA was initially prepared by heating the mix at 95°C for 5 min before cooling to room temperature. The Cas9 enzyme along with the rest of the injection mix components were then added to the guide RNA complex and incubated at room temperature for 5 minutes to assemble the RNP complexes. Microinjections were performed in *B. tryoni* Ourimbah laboratory strain embryos that were collected over 1 hour time period and prepared for injection as previously described^22^. Injections were performed under paraffin oil using borosilicate capillary needles (#30-0038, Harvard Apparatus) drawn out on a Sutter P-87 flaming/brown micropipette puller and connected to an air-filled 20 ml syringe, a manual MM-3 micromanipulator (Narishige) and a CKX31-inverted microscope (Olympus). Microscope slides with the injected embryos were placed on agar in a Petri dish that was then placed in a vented container containing moist paper towels at 25°C (± 2°C). Hatched first instar larvae were removed from the oil and transferred to larval food. Individual G_0_ flies were crossed to six virgin flies from the Ourimbah laboratory strain and eggs were collected overnight for two consecutive weeks. G_1_ flies were then allowed to mate *inter se* and eggs were collected in the same manner. G_2_ pupae were then analyzed phenotypically and separated according to color of pupae (brown, mosaic or white).

For *C. capitata* CRISPR/Cas9 gene editing, lyophilized Cas9 protein (PNA Bio Inc, CP01) was reconstituted to a stock concentration of 1 μg/μl in 20 mM Hepes, 150 mM KCl, 2% sucrose and 1 mM DTT (pH 7.5) and stored at −80°C until use. A guide RNA (gRNA_MFS), targeting the third CDS exon of *CcMFS* was designed and tested for potential off target effects using Geneious Prime^52^ and the *C. capitata* genome annotation Ccap2.1. *In silico* target site analysis predicted an on-target activity score of 0.615 (scores are between 0 and 1; the higher the score the higher the expected activity^66^ and zero off-targets sites in the medfly genome. Guide RNA was synthesized by *in vitro* transcription of linear double-stranded DNA template as previously described^67^ using primers P_1753 and P_369 (Supplementary Table 4), Q5 HF polymerase, and a Bio-Rad C1000 Touch thermal cycler [98°C, 30 s; 35 cycles of (98°C, 10 s; 58°C, 20 s; 72°C, 20 s); 72°C, 2 min] and HiScribe™ T7 High Yield RNA Synthesis. Injection mixes for microinjection of embryos contained 360 ng/μl Cas9 protein (1 μg/μl, dissolved in its formulation buffer), 200 ng/μl gRNA_MFS and an end-concentration of 300 mM KCl according to previous studies^67, 68^. The mix was freshly prepared on ice followed by an incubation step for 10 min at 37°C to allow pre-assembly of gRNA-Cas9 ribonucleoprotein complexes and stored on ice prior to injections. Microinjections were conducted in wild type EgII *C. capitata* embryos. Eggs were collected over a 30-40 min time period and prepared for injection as previously described^67^. Injections were performed using siliconized quartz glass needles (Q100-70-7.5; LOT171381; Science Products, Germany), drawn out on a Sutter P-2000 laser-based micropipette puller. The injection station consisted of a manual micromanipulator (MN-151, Narishige), an Eppendorf FemtoJet 4i microinjector, and an Olympus SZX2-TTR microscope (SDF PLAPO 1xPF objective). The microscope slide with the injected embryos was placed in a Petri dish containing moist tissue paper in an oxygen chamber (max. 2 psi). Hatched first instar larvae were transferred from the oil to larval food.

As complementation assay, reciprocal crosses between surviving G_0_ adults and virgin *white pupae* strain #1402_22m1B (*pBac_fa_attP-TREhs43-Cctra-I-hid^Ala5^-SV40_a_PUb-nls-EGFP-SV40*) (*wp*^−(nat)^) ^24^ were set up either single paired (six cages) or in groups of seven to ten flies (seven cages). Eggs were collected three times every 1-2 days. Progeny (G_1_) exhibiting the *white pupae* phenotype (*wp*^−(nat)|-(CRISPR)^) were assayed via non-lethal genotyping and sorted according to mutation genotype (see Fig. 4). Genotypes ‘A-H’ were group-backcrossed into WT EgII (*wp*^+|+^), genotype ‘C’ siblings massed crossed. Eggs were collected 4 times every 1-2 days. Generation G_2_ flies were analyzed via PCR using three primers, specific for *wp*^+^ and *wp*^−(CRISPR)^ or *wp*^−(nat)^ allele size respectively. Offspring of outcross cages showed brown pupae phenotype and either *wp*^+|− (nat)^ or *wp*^+|−(CRISPR)^ genotype. In order to make mutations A, D, and H homozygous, 40 flies (25 females, 15 males) were genotyped each, and *wp*^+|−(CRISPR)^ positive flies were inbred (mutation A: 15 females, 7 males, mutation D: 12 females, 7 males, mutation H: 11 females, 8 males). G3 showing white pupae phenotype was homozygous for *wp*^−(CRISPR)^ mutations A, D, or H, respectively and was used to establish lines. Inbreeding of mutation C *wp*^−(nat)|-(CRISPR)^ flies produced only white pupae offspring, based on either *wp*^−(nat)|-(nat)^, *wp*^−(nat)|-(CRISPR)^ or *wp*^−(CRISPR)|-(CRISPR)^. 94 flies (46 females, 48 males) were genotyped, homozygous *wp*^−^(^CRISPR)^ were inbred to establish a line (13 females, 8 males).

### Molecular analyses of *wp* mutants and mosaics

In *B. tryoni*, genomic DNA was isolated for genotyping from G_2_ pupae using the DNeasy Blood and Tissue Kit (Qiagen). PCR amplicons spanning both BtMFS guide recognition sites were generated using Q5 polymerase (NEB) with primers BtMFS_5primeF and BtMFS_exon2R (Supplementary Table 4). Products were purified using MinElute PCR Purification Kit (Qiagen), ligated into pGEM-t-easy vector (Promega) and transformed into DH5α cells. Plasmids were purified with Wizard *Plus* SV Minipreps (Promega) and sequenced.

In *C. capitata*, non-lethal genotyping was performed to identify parental genotypes before setting up crosses. Therefore, genomic DNA was extracted from single legs of G_1_ and G_2_ flies following a protocol established by ^69^. Single legs of anesthetized flies were cut at the proximal femur using scissors. Legs were then homogenized by ceramic beads and 50 μl buffer (10 mM Tris-Cl, pH 8.2, 1 mM EDTA, 25 mM NaCl) for 15 s (6 m/s) using a FastPrep-24^TM^ 5G homogenizer. Then, 30 μl buffer and 1.7 μl proteinase-K (2.5 U/mg) were added, incubated for 1 h at 37°C and the reaction stopped by 4 min at 98°C. The reaction mix was cooled down on ice and used for PCR. For G_1_ flies, PCR on *wp* was performed in a 25 μl reaction volume using the Dream*Taq* polymerase, primer pair P_1643 and P_1644 (Supplementary Table 4), and 3.75 μl reaction mix. Different amplicon sizes are expected for brown (*wp*^+^ and *wp*^−(CRISPR)^) and white pupae (*wp*^−(nat)^) alleles, 724 bp and 8,872 bp in size, respectively. The *wp*^−(nat)^ amplicon was excluded via PCR settings [95°C, 3 min; 35 cycles of (95°C, 30 s; 56°C, 30 s; 72°C, 1 min); 72°C, 5 min]. The 724 bp PCR product was verified by agarose electrophoresis and purified from the PCR reaction using the DNA Clean & Concentrator^TM^-5 kit. PCR products were sequenced using primer P_1644 and sequences analyzed using the Geneious Prime Software Package (6). In generation G_2_, flies were analyzed using multiplex PCR with primers P_1657, P_1643, and P_1644 (Supplementary Table 4), to distinguish between the *wp*^−(nat)^ (457 bp; P_1643 and P_1657), and *wp*−^(CRISPR)^ alleles (724 bp; P_1643 and P1644) using the PCR protocol [95°C, 3 min; 35 cycles of (95°C, 30 s; 56°C, 30 s; 72°C, 1 min); 72°C, 5 min]).

### Image acquisition

Images of *B. tryoni* pupae were taken with an Olympus SZXI6 microscope, Olympus DP74 camera and Olympus LF-PS2 light source using the Olympus stream basic 2.3.3 software. Images of *C. capitata* pupae were taken with a Keyence digital microscope VHX-5000. Image processing was conducted with Adobe Photoshop CS5.1 software to apply moderate changes to image brightness and contrast. Changes were applied equally across the entire image and throughout all images.

## Ethics approval and consent to participate

Not applicable

## Consent for publication

Not applicable

## Availability of data and material

All data generated or analyzed are included in this article.

## Acknowledgements

This study was benefitted from discussions at International Atomic Energy Agency funded meetings for the Coordinated Research Projects (CRPs) and particularly the CRP on “Generic approach for the development of genetic sexing strains for SIT applications”. The authors also wish to thank Tanja Rehling, Jakob Martin and Johanna Rühl for technical assistance and Germano Sollazzo for helping with injections and primers design (Justus-Liebig University Gießen and Insect Pest Control Laboratory); Elena Isabel Cancio Martinez, Thilakasiri Dammalage, Sohel Ahmad and Gülizar Pillwax for insect rearing (Insect Pest Control Laboratory), Shu-Huang Chen (McGill University) for technical assistance with Nanopore library preparations; and Arjen van’t Hof (University of Liverpool) for constructing libraries from micro-dissected chromosomes.

